# Structural basis for BCL7B-mediated ncBAF-nucleosome engagement

**DOI:** 10.1101/2025.11.20.689410

**Authors:** Fahui Sun, Binqian Zou, He Li, Chongshen Xu, Qiaohong Luo, Chi Wang, Pengqi Xu, Duanqing Pei, Jiekai Chen, Dajiang Qin, Ying Zhang, Jun He

## Abstract

The mammalian SWI/SNF family of chromatin remodelers comprises BRG1/BRM-associated factor (cBAF), Polybromo-associated BAF (PBAF), and non-canonical BAF (ncBAF) complexes, which slide and disassemble nucleosomes to regulate gene expression and chromatin structure dependent on ATP hydrolysis energy. While the chromatin engagement mechanisms of cBAF and PBAF have been structurally resolved, the molecular architecture governing ncBAF interaction with chromatin remains elusive. In this study, by integrating cryo-electron microscopy, biochemical assays and cross-linking mass spectrometry, we resolved the conformational transition of ncBAF-nucleosome complexes from nucleotide-free to nucleotide-bound states. Our analyses establish BCL7 proteins as dynamic molecular tethers connecting the ARP module to the nucleosomal acidic patch and demonstrate that BCL7B promotes ncBAF-mediated nucleosome remodeling, with BRG1-catalyzed ATP hydrolysis triggering conformational changes that modulate BCL7-mediated histone association. Structurally and biochemically, we further demonstrate that β-actin within the BCL7-containing ARP module retains ATP hydrolysis activity, rendering its exposed pointed end structurally compatible with incorporation into the barbed end of nuclear actin filaments, which provides a potential molecular basis for coordinating nuclear actin networks with chromatin remodeling. Collectively, our findings unravel a dynamic role of BCL7 in regulating ncBAF-mediated chromatin remodeling and establish a distinct chromatin engagement mode of ncBAF from that of cBAF/PBAF.

## Introduction

Chromatin remodeling complexes serve as essential regulators of gene expression and chromatin dynamics in response to cellular signals (1). Among these complexes, the Switch-Sucrose Non-Fermentable (SWI/SNF) family of chromatin remodelers plays a critical role in ATP-dependent chromatin remodeling through the concerted action of various factors (2). This multi-subunit complex family shares a common catalytic subunit (SMARCA2/4, also known as BRM/BRG1) and is categorized into three subtypes based on their accessory components: canonical BRG1/BRM-associated factor (cBAF), Polybromo-associated BAF (PBAF), and non-canonical BAF (ncBAF) complexes (3). While the first two subtypes have conserved counterparts in yeast, namely Swi/Snf and RSC complexes, ncBAF appears to be specific to higher organisms (4). These SWI/SNF complexes share a core set of subunits but are distinguished by unique components that confer each complex with distinct functional properties. Notably, ncBAF lacks several subunits present in cBAF and PBAF, such as SMARCB1, SMARCE1, SMARCD2/3, and SMACC2, but incorporates unique subunits like BRD9 and GLTSCR1/GLTSCR1L (2). Given the essential roles of SWI/SNF complexes in pluripotency, cancer, and neurodevelopment, several subunits of these complexes have been identified as promising therapeutic targets by perturbing SWI/SNF functions (2). In particular, ncBAF proved to be a synthetic lethal target in synovial sarcoma and malignant rhabdoid tumors carrying SMARCB1 mutations (5).

The architecture of the SWI/SNF complex has been extensively studied in both yeast and mammals (6–11). In humans, a catalytic subunit scaffolds these complexes, either BRG1 or BRM, around which auxiliary subunits sequentially assemble to form a modular architecture that includes the base module, the actin-related protein (ARP) module, and the ATPase module (2). Cryo-EM studies have provided detailed insights into the interaction of cBAF and PBAF complexes with nucleosomes, revealing that both complexes similarly engage nucleosomes, with the ATPase and base modules sandwiching the nucleosome (10,11). Specifically, on one side of the nucleosome, BRG1 is positioned at the superhelical location +2 (SHL+2) of nucleosomal DNA, interacting with both the acidic patch and the histone H4 tail. On the opposite side, SMARCB1 offers a helix that engages the second acidic patch. However, to date, no studies have elucidated the ncBAF-nucleosome structure, leaving unclear how ncBAF interacts with the chromatin.

The yeast ARP module has been implicated in facilitating chromatin remodeling activity of the SWI/SNF complexes (12). To date, however, direct physical interactions between the ARP module and the nucleosome have not been delineated in either yeast or human SWI/SNF-nucleosome complexes (6,8–11). In the yeast ARP module, the subunit Rtt102 employs a β-sheet fold that contributes to the assembly of Arp7, Arp9, and the Snf2 HSA domain (13–15). An equivalent β-sheet fold has been observed in human chromatin remodeling complexes SRCAP, TIP60, and yeast acetyltransferase NuA4 complex (16–20). These folds arise from non-homologous subunits, highlighting their evolutionary conservation in different chromatin-associated complexes across species. Despite the critical role of the β-sheet fold, homologs or any counterpart of Rtt102 have yet to be identified in human SWI/SNF complexes, raising the question of how the ARP module is intrinsically assembled in mammals. Recent studies suggest that BCL7 family proteins (including BCL7A, BCL7B, and BCL7C) interact with the HSA domain of BRG1, and benefit chromatin remodeling and gene transcription (21). This finding implies that BCL7 proteins might functionally replace Rtt102 to complete the ARP module in mammals. While this emerging evidence sheds light on the potential role of BCL7 proteins in human SWI/SNF complexes, the precise structural mechanism of these interactions remains poorly understood, warranting further investigation to elucidate this mechanism in chromatin remodeling.

In this study, we determined the ncBAF-nucleosome structures both in the presence and absence of ATP analogs, revealing a distinct interaction mode compared to the previously characterized cBAF/PBAF-nucleosome complexes. Our findings demonstrate a significant conformational shift triggered by BRG1-catalyzed ATP hydrolysis, which modulates the function of BCL7B as a structural bridge between the ARP module and the nucleosomal acidic patch. In contrast to cBAF and PBAF, the base module of ncBAF does not directly engage with the nucleosomal acidic patch, suggesting a unique mechanism of chromatin recognition. Furthermore, our high-resolution structures of the BCL7-containing ARP module elucidated the mechanistic basis of BCL7 incorporation into the ARP module. Unexpectedly, the structures revealed that the β-actin subunit is capable of binding either ATP or ADP, indicating an intrinsic ATPase activity that was further confirmed biochemically. Moreover, the architecture of the ARP module shows that the exposed pointed end of β-actin is structurally compatible with incorporation into the barbed end of an actin filament. This architectural feature suggests a potential interaction mode through which the SWI/SNF complexes interact with nuclear actin filaments. Collectively, our findings offer a novel structural framework for how mammalian SWI/SNF complexes engage with nucleosomes and highlight the pivotal role of BCL7 in regulating ncBAF-mediated chromatin remodeling, distinguishing it from the mechanisms operating in cBAF and PBAF.

## Materials and methods

### Protein complexes expression and purification

The nine full-length coding sequences of human ncBAF subunits (BRG1, GLTSCR1, SMARCC1, BRD9, BAF60A, ACTL6A, β-actin, SS18, BCL7B) were subcloned into pcDNA3.4 vectors, respectively. A 2×Strep tag II was fused to the C-terminus of GLTSCR1 to facilitate the complex purification. All the expression constructs were co-transfected into the HEK293F suspension cells using polyethylenimine (Polysciences) at a cell density of 2×10^6^/ml. After culturing for 3 days in conditions of 37℃ and 5% carbon dioxide, cells were collected by centrifugation at 1000×g. The cell pellets were flash-frozen in liquid nitrogen and stored at -80℃, or resuspended in lysis buffer (50 mM HEPES, pH 8.0, 200 mM NaCl, 100 mM Arginine and Glutamate, 0.1% NP-40, 0.5 mM EDTA, 2 mM MgCl_2_, 2 mM DTT, DNase I and protease inhibitor cocktail) for protein purification at 4℃. After ultrasonic disruption, the cell lysate was centrifuged at 20000 rpm for 1 hour. The supernatant was loaded into a Strep-tactin column overnight and then thoroughly washed the column using washing buffer (25 mM HEPES, pH 8.0, 250 mM NaCl, 50 mM Arginine and Glutamate, 0.02% NP-40, 2 mM MgCl_2_, 2 mM DTT, 1 mM ATP, and 0.5 mM PMSF). The protein complex was then eluted using elution buffer (25 mM HEPES, pH 8.0, 100 mM NaCl, 25 mM Arginine and Glutamate, 1 mM MgCl_2_, 2 mM DTT, and 5 mM D-desthiobiotin). To remove nucleic acid contaminants, we further processed the eluate using an anion exchange column (MonoQ 5/50 GL column, GE Healthcare). The fractions corresponding to the ncBAF complex were combined and concentrated to 6-10 mg/ml using preservation buffer (25 mM HEPES, pH 7.5, 150 mM NaCl, 1 mM MgCl_2_, and 2 mM DTT). The final ncBAF protein was directly used to prepare cryo-EM samples or flash-frozen in liquid nitrogen and stored at -80℃. The BCL7B N-terminal helix (residues 1-20) deleted version of the ncBAF complex was expressed and purified following the same procedure as the wild-type complex. For purification of the EGFP-fused ARP-ATPase subcomplex of ncBAF used in the biophysical assays, the N-terminal 1-462 amino acid region of BRG1 was replaced with the EGFP sequence. The expression plasmid encoding the EGFP-BRG1 fusion protein was co-transfected into HEK293F cells together with ACTL6A, β-actin and BCL7B, followd by purification using a procedure similar to that employed for the full ncBAF complex.

For purification of the BCL7B-containing ARP module, a similar protocol was followed. In brief, full-length BCL7B, ACTL6A, β-actin, and HSA domain of BRG1 (447-518 AA, with a C-terminal 2×Strep tag II) were co-transfected into HEK293F cells. After 72-hour culture, cells were harvested and ultrasonically disrupted in lysis buffer (50 mM Tris-HCl, pH 8.0, 200 mM NaCl, 1 mM EDTA, 2 mM MgCl_2_, 0.05% Triton X-100, 1 mM PMSF, 3 mM DTT, DNAse I, 1 mM ATP, and protease inhibitors). The lysate was clarified by centrifugation at 20000 rpm for 1 hour, followed by the supernatant loading onto a Strep-tactin column and thorough washing with lysis buffer. Subsequently, the eluate was further polished with an anion exchange column (MonoQ 5/50 GL column, GE Healthcare). Finally, the fractions containing the ARP module complex were collected and ultrafiltered using an exchange buffer (25 mM HEPES, pH 7.4, 50 mM NaCl, 1 mM MgCl_2_, and 2 mM DTT). The concentrated ARP module complex was utilized for cryo-EM analyses. For purification of the ARP module carrying the β-actin (Q137A) mutant, an identical purification procedure was used as for the wild-type complex.

### Preparation of nucleosomes

Three types of nucleosomes were reconstituted for different experimental purposes, following the methods modified from the previous literature (22,23). The 0N20 nucleosome, containing 147 bp of Widom 601 positioning sequence flanked by 20-bp linker DNA on one side, was used for cryo-EM structural studies of ncBAF-nucleosome complexes. Two variants of 0N0 nucleosomes, consisting of only the 147-bp Widom 601 sequence without linker DNA, were prepared for FRET-based conformational analyses: one labeled with Atto425 fluorophore at DNA ends, and the other labeled on the mutant H2B(S123C) residue using 5(6)-TAMRA C6 maleimide (AAT Bioquest, cat# 423). The 100N100 nucleosome, containing the 147-bp Widom 601 sequence flanked symmetrically by 100-bp extranucleosomal DNA on both sides, was employed in restriction enzyme accessibility assay of remodeling assays.

For the Xenopus octamer purification, four core histones were subcloned into one pRSFDuet vector in a polycistronic way, and co-expressed in Escherichia coli BL21 (DE3) cells. The harvested cell pellets were resuspended in lysis buffer (50 mM Tris-HCl, pH 8.0, 500 mM NaCl, 0.1 mM Na-EDTA, pH 7.5, 1 mM PMSF, and 3 mM DTT) for subsequent high-pressure homogenization. After centrifugation at 18000 rpm for 0.5 hour, the supernatant was directly applied to an ion-exchange column (HiTrap Heparin HP column, Cytiva) for the histone octamer enrichment. The eluted fractions containing the octamer were combined and concentrated to run a further size-exclusion chromatography (HiLoad 16/600 Superdex 200, Cytiva) using a high salt buffer (20 mM Tris-HCl, pH 8.0, 2 M NaCl, and 2 mM DTT). The mutant octamer (H2BS123C) used for the fluorescence resonance energy transfer (FRET) was expressed and purified following the same procedure as for the wild-type octamer.

The DNA chains containing Widom 601 DNA sequence (5’-CTGGAGAATCCCGGTGCCGAGGCCGCTCAATTGGTCGTAGACAGCTCTAGCACCGCTTAAACGCACGTACGCGCTGTCCCCCGCGTTTTAACCGCCAAGGGGATTACTCCCTAGTCTCCAGGCACGTGTCAGATATATACATCCTGT-3’) were amplified from a plasmid template using PCR. For FRET nucleosome assembly, PCR primers labeled with Atto425 at the 5’ end were used to fluorescently tag the DNA fragment. PCR products were purified by anion-exchange chromatography (Resource Q column, Cytiva), and the target fragment was collected by isopropanol precipitation and stored at -20 °C. Before use, the DNA was dissolved in 2 M NaCl solution, and concentration was measured by NanoDrop at 260 nm.

Nucleosomes were assembled by salt-gradient dialysis. The purified histone octamer and DNA fragment were mixed at a molar ratio of 1.2:1 and gradually dialyzed from 2 M to 250 mM salt buffer for more than 20 hours. The reconstituted nucleosomes were further dialyzed into low-salt buffer (10 mM HEPES, pH 7.5, 30 mM NaCl) for long-term storage. Nucleosome homogeneity was confirmed by native-PAGE prior to use in structural and biochemical assays.

### ncBAF-nucleosome complexes reconstruction

To assemble the ncBAF-NCP complexes in the apo state for cryo-EM analysis, the purified ncBAF (3 uM) and 0N20 nucleosome (1.5 uM) were mixed in a buffer containing 25 mM HEPES, pH 7.5, 200 mM NaCl, 1 mM MgCl_2_, and 2 mM DTT. Further, the mixture was dialyzed at 4℃ for 3 hours in a low-salt buffer (25 mM HEPES, pH 7.5, 30 mM NaCl, 1 mM MgCl_2_, and 2 mM DTT). The complexes were then cross-linked and isolated using GraFix method as previously described (24). Fractions containing ncBAF-NCP complex were collected and ultrafiltered in cross-linker quench buffer (50 mM Tris, pH 7.5, 30 mM NaCl, 1 mM MgCl_2_, and 2 mM DTT). As for the assembly of the ADP-BeFx-bound ncBAF-NCP complex, the same procedure was followed, except the buffers for dialysis and GraFix were supplemented with 1 mM ADP, 2 mM BeSO_4_, and 16 mM NaF.

### Cryo-EM grids preparation and data collection

Quantifoil R1.2/1.3 holey carbon grids (300 mesh) were glow-discharged in air for 30 seconds at 20 mA using a GloQube Plus system. Grids were subsequently mounted onto a Vitrobot Mark IV (Thermo Fisher Scientific) operated at 4°C and 100% humidity. A 4 μL aliquot of ncBAF–NCP complex, either wild-type or deletion mutant, at a concentration of 1 mg/mL, was applied to each grid. Grids were blotted using a force setting of 3 for 2 seconds and plunge-frozen in liquid ethane cooled by liquid nitrogen. Grids containing ARP module complexes, in either the apo form or incubated overnight with 1 mM ADP, were prepared following the same protocol.

Cryo-EM data were acquired using a Titan Krios G4 transmission electron microscope (Thermo Fisher Scientific) equipped with a Falcon 4 direct electron detector and a SelectrisX energy filter set to a 10 eV slit width. Automated data collection was performed in electron event representation (EER) mode using the EPU software. For the apo ncBAF–NCP sample, 25,525 movies were recorded at a dose rate of ∼5.51 e⁻/Å²/s with a nominal pixel size of 0.93 Å at the specimen level. Imaging was conducted with a defocus range of -0.8 to -2.4 μm and a total exposure time of 7.44 seconds, resulting in a cumulative dose of ∼50 e⁻/Å². For the ADP-BeFx-bound ncBAF-NCP, apo ARP, and ADP-soaked ARP samples, 11,737, 4,972, and 8,190 movies were collected, respectively, under similar conditions. These datasets were acquired at a dose rate of ∼5.37 e⁻/Å²/s and a pixel size of 0.73 Å, with identical defocus settings and a total exposure time of 4.68 seconds, yielding a total dose of ∼50 e⁻/Å².

### Image processing

Image stacks were processed using CryoSPARC software (v 4.4.0) (25). The EER movies were subjected to patch motion correction and patch CTF estimation. Particles were picked by Blob picker and selected through multiple rounds of 2D classification. By using 2D classification and the stochastic gradient descent algorithm, several initial models were obtained for subsequent processing. The particles in clearly defined 2D classes were then subjected to several rounds of heterogeneous refinement to isolate unique conformational states and eliminate particles in poorly defined classes. For the apo ncBAF-NCP sample, global refinement maps were performed with the CryoSPARC non-uniform refinement, and subtracted particles with the mask of the NCP and ARP module for local refinement. After the ARP module focused on no-alignment 3D classification, the final map was produced through CryoSPARC local refinement with masks including only the NCP or ARP module. Composite maps were generated with Chimera X (v.1.7) (26). For the ADP-BeFx-bound ncBAF-NCP dataset processing, a heterogeneous analysis was performed in cryoDRGN using principal component analysis method (27). For ATP-ARP and ADP-ARP datasets, final maps were performed with the CryoSPARC non-uniform refinement, which was filtered by heterogeneous refinement. The local resolution map was generated using RELION (28) and visualized with UCSF ChimeraX. All reported resolutions are based on the gold-standard Fourier shell correlation (FSC) 0.143 criterion.

### Model building

To facilitate module building and analysis, map sharpening was performed with CryoSPARC, deepEMhancer (29), EMReady v2.0 (30) and locSpiral (31). The cBAF-nucleosome structure was used as an initial structural template for modeling (10). The model was docked into the cryo-EM density maps using UCSF ChimeraX. Initial fitted models were manually adjusted in COOT (32), followed by multiple rounds of real-space refinement using Phenix (v.1.20.1-4487) (33) and Namdinator (34). Validation of cryo-EM maps and models was performed with Phenix. Protein interaction analysis was conducted using PLIP (35) and PISA (36).. The structure figures were generated using UCSF ChimeraX.

### FRET assay

Fluorescence resonance energy transfer (FRET) assays were performed in a buffer containing 20 mM Tris-HCl (pH 7.5), 80 mM NaCl, 1 mM MgCl_2_, 5% (v/v) glycerol, 0.01% (v/v) NP-40, 0.1 mg/ml BSA, and 2 mM DTT. For ATP analog-bound conditions, 1 mM ADP, 2 mM BeSO₄, and 16 mM NaF were added to generate the ADP-BeFx complex. The fluorophore-labelled nucleosome core particle (NCP) and EGFP-fused ATPase-ARP subcomplex were each used at 100 nM (molar ratio 1:1). Samples were incubated at room temperature at least for 30 min, and fluorescence signals were measured using FlexStation 3 multimode plate reader. The excitation wavelengths for the Atto425 and EGFP fluorophores were 436 nm and 488 nm, respectively. The emission detection cutoffs were set at 480 nm for Atto425, 520 nm for EGFP, and 580 nm for TAMRA. Each condition was measured in five independent replicates. The background signal from the corresponding buffer was subtracted prior to data analysis. FRET efficiency was calculated as:

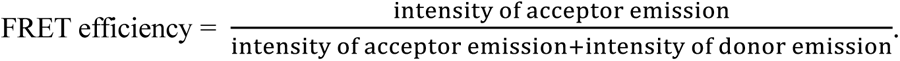

The resulting efficiencies were reported as mean ± SEM.

### Cross-linking mass spectrometry analysis

For cross-linking mass spectrometry, wild-type ncBAF (1mg/ml) was mixed with 0N20 nucleosomes at a 2:1 molar ratio in buffer containing 25 mM HEPES, pH 7.5, 150 mM NaCl, 2 mM MgCl_2_, and 2 mM DTT, either in the presence or absence of ADP-BeFx. Samples were cross-linked on ice with 1 mM BS3 (bis(sulfosuccinimidyl)suberate) for 50 minutes, and the reaction was quenched with 60 mM Tris-HCl. Cross-linked complexes were submitted to the National Facility for Protein Science (Shanghai) for mass spectrometry analyses, and the XL-MS data were visualized using xiView (37).

### Fluorescence polarization binding assay

Fluorescein isothiocyanate (FITC) was conjugated to the 5′ end of one primer used to amplify the Widom 601 DNA segment. FITC-labeled nucleosomes were then reconstituted as described above. Fluorescence polarization binding assays were carried out by incubating 20 nM FITC-nucleosome with two-fold serial dilutions of ncBAF complexes (wild-type or deletion mutant), spanning 1 µM to 0.00098 µM. The binding buffer consisted of 20 mM Tris-HCl (pH 7.5), 80 mM NaCl, 5% glycerol, 0.01% NP-40, 0.1 mg/mL BSA, and 2 mM DTT. Reactions were incubated at room temperature for 1 hour, then polarization was measured on an EnVision multimode plate reader with the gain set to 27. Each condition was assayed in triplicate, and a control containing only 20 nM FITC-nucleosome was included. The background (control value) was subtracted from all measurements before analysis. Data were fitted to a specific-binding model with Hill slope using GraphPad Prism 9, and dissociation constants are reported as mean ± SEM.

### Remodeling assay

The nucleosome remodeling assay was adapted from a previous study (38) and performed at 30°C. Reactions contained 50 nM ncBAF complex and 50 nM nucleosomes with 100-bp flanking DNA, in buffer composed of 25 mM HEPES (pH 7.4), 50 mM KCl, 2 mM MgCl₂, 4 U/µl MfeI (NEB), 4 U/µl PmlI (NEB), 0.1% NP-40, 2% glycerol, and 2 mM DTT. After a 5-minute preincubation, ATP was added to a final concentration of 4 mM to initiate remodeling. Five-microliter aliquots were collected at the indicated time points and quenched with 2× stop buffer (25 mM HEPES, pH 7.4, 50 mM KCl, 2% glycerol, 2 mM DTT, 1% SDS, 100 mM EDTA, and 2 mg/ml proteinase K (Beyotime)). Samples were deproteinized at 50°C for 30 min and resolved on 5% native-PAGE gel for 60 min at 100 V. Gels were stained with GelRed (Vazyme) and imaged. Each remodeling assay was performed in three independent technical replicates. The intensity of the uncleaved 347-bp DNA band, quantified by integrated density, was used to calculate the remodeled fraction relative to the value at 0 min. Data (mean ± SEM) were fitted to one-phase decay curves using GraphPad Prism 9.

### ATPase hydrolysis assay

ATPase hydrolysis activity was measured using the ADP-Glo^TM^ Kinase Assay Kit (Promega, V6930) according to the manufacturer’s instructions. Each reaction contained 100 nM of the indicated protein or complex and was incubated for 30 min at room temperature. The signal from the buffer-only control (25 mM HEPES, pH7.4, 100 mM NaCl, 2 mM MgCl2, 1 μM ATP, 0.5 mM DTT) was used as background and subtracted from all measurements. Each condition was assayed in independent triplicates, and the final hydrolysis rates were reported as the mean ± SEM, plotted using GraphPad Prism 9.

## Results

### BRG1 ATP hydrolysis induces ARP module reorientation in ncBAF-NCP structures

To investigate the structural architecture of the nucleosome-bound ncBAF complex, we recombinantly expressed and purified *homo sapiens* ncBAF complex in HEK293F cells (Figure S1). Using 167-bp DNA fragments composed of a 20-bp linker followed by the canonical Widom 601 sequence, we reconstituted nucleosomes that were subsequently complexed with ncBAF under gradient fixation (GraFix) conditions (24). This approach yielded ncBAF-nucleosome complexes in the presence and absence of ADP-BeFx (a non-hydrolyzable ATP analog). Both samples were then subjected to cryo-EM single-particle analyses, resulting in 3D maps of the ncBAF-nucleosome complex at the apo and ADP-BeFx-bound states, with overall resolutions of 3.7 and 3.05 Å, respectively (Figure 1A, B, Figure S2, S3, Table 1).

**Figure 1.**
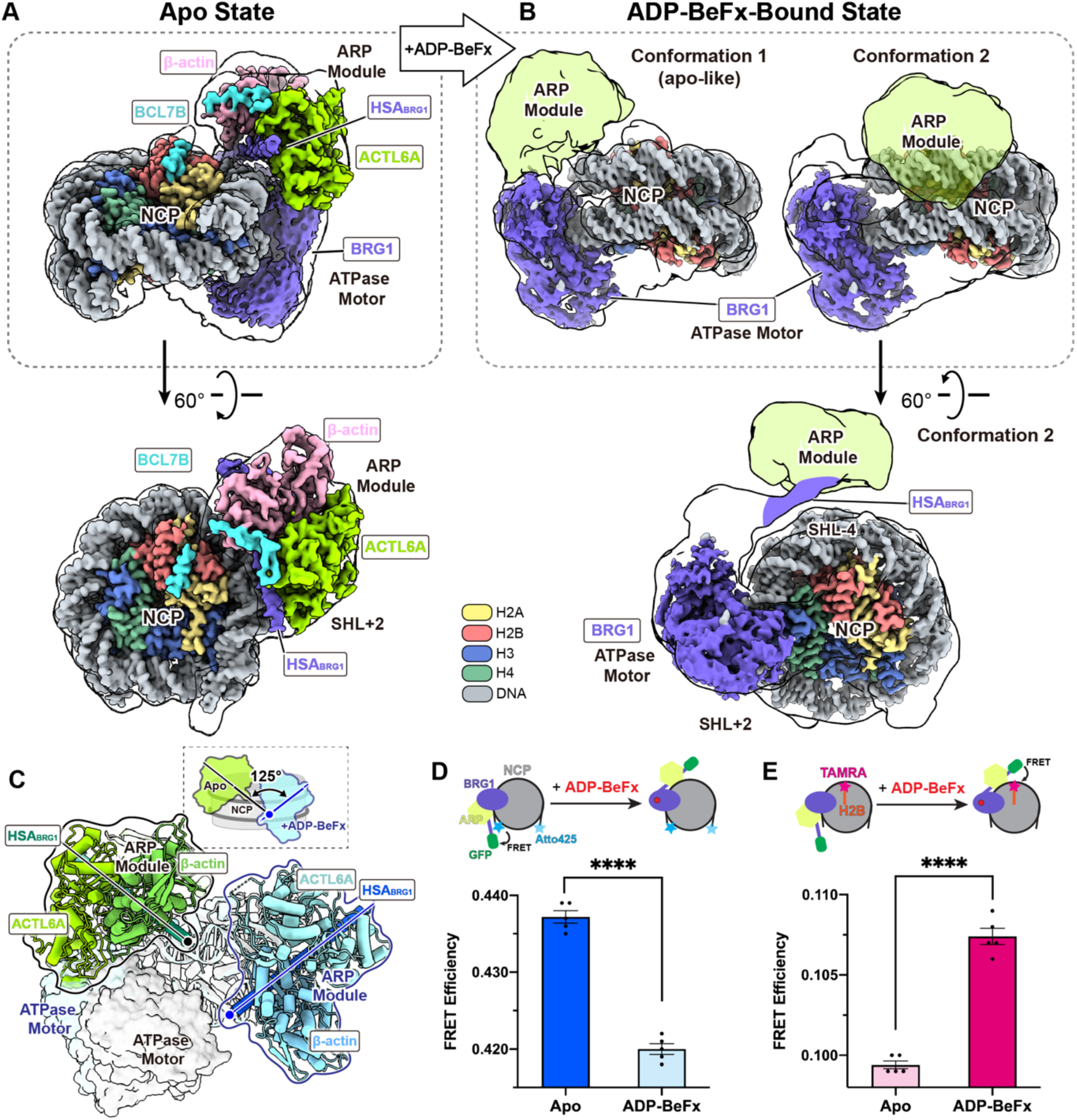
Cryo-EM structures of the human ncBAF-NCP complexes in the nucleotide-free and -bound states. (**A**) Composite cryo-EM map of the ncBAF-nucleosome complex in the apo state, colored as indicated and shown in two orientations. (**B**) Cryo-EM maps of two distinct conformations identified in the ADP-BeFx-bound ncBAF-nucleosome complex, colored as indicated as in panel (A). The 3.7-Å consensus map from Figure S2A and the two conformational maps from Figure S3C are displayed as transparent overlays in panels (A) and (B). (**C**) Superposition of the nucleotide-free and nucleotide-bound (conformation 2) ncBAF–nucleosome complexes aligned on the nucleosome. The ARP modules are shown as tube models (green for apo and blue for ADP–BeFx–bound), with the cryo-EM densities of the ATPase motor modules displayed. Upon ADP–BeFx binding, the long helix of the BRG1 HSA domain rotates by approximately 125°, repositioning the ARP module closer to the nucleosomal DNA at the SHL-4 site. (**D** and **E**) Fluorescence resonance energy transfer (FRET) analyses of ATP analog–induced conformational changes in the ncBAF–nucleosome complex. (**D**) Schematic of the FRET assay using Atto425-labeled nucleosomes (Atto425–NCP) and an EGFP fusion to the N terminus of BRG1 (Δ1–462) to monitor the distance between the ARP module and the nucleosomal DNA ends. The bar graph shows a marked decrease in FRET efficiency upon ADP–BeFx binding, indicating that the ARP module moves away from the nucleosomal DNA ends. (**E**) Schematic of the FRET assay using TAMRA-labeled nucleosomes (TAMRA–NCP) with the fluorophore attached to H2B S123C near the SHL-4 region. The bar graph shows a statistically significant increase in FRET efficiency in the ADP–BeFx–bound state. FRET data are presented as mean ± standard error of the mean (SEM) from five technical replicates. Statistical significance was determined using a two-tailed Student’s *t*-test (****p < 0.0001).

The cryo-EM map of the apo ncBAF-NCP structure suggests that the ATPase motor of ncBAF engages the nucleosome core particle (NCP) at the SHL+2 position (Figure 1A), consistent with previous studies of ATPase remodeling complexes in the same family (6–11). Apart from the ATPase motor and nucleosome, local refinement of the ARP module density yielded a reconstruction of 4.13 Å resolution, which enabled the unambiguous assignment of the HSA helix, β-actin, and ACTL6A subunits of the ARP module (Figure 1A, Figure S2A). Of particular interest, the local refinement analysis revealed an area of unknown density adjacent to ACTL6A and β-actin, a feature that will be discussed in greater detail subsequently. This observation is largely similar to the previously identified yet uncharacterized density in the ARP module of the PBAF-NCP complex (39).

In previous structural studies of the ATPase remodeling complex, ADP-BeFx has typically been utilized to stabilize an ATP-bound but non-hydrolyzed state (7,8). In our study, initial cryo-EM data processing of the ADP-BeFx-bound ncBAF-NCP sample yielded a density map of 3.05 Å resolution, clearly revealing the two-lobe feature of the ATPase motor (Figure S3A). Surprisingly, we no longer observed the density for the ARP module (Figure S3A). To characterize the ARP module, particles from the 3.05-Å map were subjected to a heterogeneous analysis using cryoDRGN (27), which revealed a density transition trajectory with a putative ARP density waxing and waning between two distinct positions (Figure S3B). This result prompted us to do extensive classification of particles use a mask focused on the two positions. Reprocessing revealed two conformations: conformation **1** (∼19% particles) resembling the apo ncBAF-NCP structure, and conformation **2** (∼81% particles), exhibiting a substantial shift of the ARP module toward the nucleosomal DNA at SHL-4 (Figure 1B, Figure S3C). Therefore, we determined that the binding of ADP-BeFx to BRG1 triggered a substantial repositioning of the HSA helix, along with the entire ARP module, resulting in a significant rotation of approximately 125 degrees relative to its position in the apo state (Figure 1C). This finding is consistent with a previous structural study in which the yeast RSC subcomplex (only containing Sth1, Arp7, Arp9, Rtt102 and lacking the base module) bound to nucleosomes adopted a conformation similar to our ncBAF-nucleosome complex in the presence of ADP-BeFx (38). Together, these observations indicate a conserved mechanism from yeast to humans, whereby the catalytic subunits of SWI/SNF complexes undergo a substantial conformational shift orchestrated through the ATPase motor driven by the ATP hydrolysis cycle.

In addition, fluorescence resonance energy transfer (FRET) assays were conducted to validate the conformational shift of ARP module. Based on structural analysis, an ncBAF subcomplex lacking the base module and carrying an EGFP fused to the N-terminus of the BRG1 HSA domain was purified for FRET assays, close mirroring the yeast RSC subcomplex described above. The EGFP in this ncBAF subcomplex was designed to form an FRET pair with fluorophores attached either to the nucleosomal DNA end under apo conditions or to H2B C-terminus on the nucleosome in the ADP-BeFx-bound state. Upon ADP-BeFx induction, the FRET efficiency showed statistically significant changes consistent with the expected conformational shift (Figure 1D, E). The relatively small amplitude of these changes likely reflects conformational heterogeneity or distances beyond the optimal FRET range.

Despite extensive efforts, the base module of ncBAF, comprising subunits SMARCC1, SMARCD1, BRD9, GLTSCR1, and SS18, remains unresolved in our cryo-EM maps. The raw particle dimensions, approximately 21 nm, closely correspond to the full ncBAF-NCP complex and are consistent with the expected size of cBAF/PBAF-NCP assemblies (Figure S4). The absence of the base module in our cryo-EM maps of the ncBAF-NCP complex strongly suggests its structural flexibility. Unlike in cBAF/PBAF-NCP structures, where SMARCB1 engages the nucleosomal acidic patch, no direct interactions with the histone core stabilize the base module in ncBAF-NCP structures. Notably, SMARCB1 appears to stabilize the base module even in the presence of ATP-analog, suggesting that the conformational shift of BRG1 is relatively modest and can be accommodated, consistent with our structural heterogeneity analysis. To further confirm this insight, we performed cross-linking mass spectrometry (XL-MS) analyses to assess the spatial proximity of subunits within the ncBAF-NCP complexes (Figure S5A, B). Our analyses, together with previously published XL-MS data of cBAF/PBAF-NCP complexes, allowed us to compile a comprehensive list of SWI/SNF subunits that are cross-linked with histones (Figure S5C). For the cBAF/PBAF-NCP complexes, whether bound to ADP-BeFx or in the apo state, 3-7 distinct cross-linked subunit pairs were identified between histones and base module components (e.g., SMARCB1, SMARCC2, PBRM1, ARID2, PHF10 in PBAF and SMARCB1, SMARCC2, SMARCC1, ARID1A, DPF2 in cBAF). In contrast, only a single cross-linked subunit pair (H3-SMARCC1) was detected in the apo state of the ncBAF-NCP complex, highlighting a unique interaction mode of ncBAF with the nucleosome. Independent of ATP-analog conditions, cross-links between BRG1 and ARP subunits in our cross-linking result predominantly map to the HSA domain and ATPase lobe 1 of BRG1, consistent with their interactions and spatial proximity observed in our cryo-EM maps. Notably, in the presence of ADP-BeFx, fewer cross-linked subunit pairs between histones and the ARP/base module of ncBAF were detected (Figure S5C), reflecting that the ARP module is positioned farther from the histones following the conformational shift in BRG1.

Taken together, these findings suggest that, unlike cBAF and PBAF, the base module of ncBAF is not effectively stabilized by the mono-nucleosome substrate and is undetectable in our cryo-EM maps. Specifically, the absence of a SMARCB1-like component in ncBAF, which contributes to nucleosome interactions in cBAF and PBAF, results in substantial flexibility of the base module in the ncBAF-NCP complex. Further exploration is needed to determine whether the ncBAF base module exhibits enhanced recognition or interaction with other types of nucleosome substrates, such as di-nucleosomes or subnucleosomes, as these substrates have yet to be investigated.

### BCL7B bridges the ARP module and histones in a BRG1-ATPase dependent manner to promote nucleosome remodeling

The SnAC domain of BRG1 (residues 1332-1390) extends from ATPase lobe 2 and engages the nucleosomal acidic patch in the PBAF-NCP structure through a characteristic arginine anchor (Arg1372) (Figure S6A). In our ncBAF–NCP maps, similar densities project from the ATPase motor toward the acidic patch (Figure 1A, B), and arginine-anchor-like densities become evident at higher contour levels (Figure S6A). These observations indicate that BRG1 engages the nucleosome in a similar manner across SWI/SNF remodelers. Given this shared mode of BRG1 engagement, and considering the increased flexibility of the ARP module in the nucleotide-bound state of the ncBAF-nucleosome complex, we next sought to understand how the ARP module is stabilized in the apo state. We noticed that the ARP module of ncBAF was positioned closer to the nucleosome compared to the cBAF or PBAF (Figure S6B), with some density extending from the ARP region towards the histones and finally seating on the nucleosomal acidic patch (Figure 2A). Subsequent local refinement of the nucleosome, including this additional density, revealed this density as a helix (Figure 2B and S8B), which we de novo modeled as the conserved N-terminal region (residues 2-18) of BCL7B protein (Figure 2B, Figure S8A-C). From the nucleosome center to the periphery, BCL7B residues Arg4, Arg7, Arg11, and Lys15 collectively form a basic surface that interacts with the acidic patch of the nucleosome (Figure 2B, Figure S8C). Notably, Arg4 serves as a characteristic arginine anchor, a hallmark feature frequently observed in chromatin-associated proteins, playing a pivotal role in nucleosome recognition and binding (40). As described above, the binding of ADP-BeFx to BRG1 results in a significant conformational change that moves the ARP module away from the nucleosome. This alteration in conformation is anticipated to facilitate the release of BCL7B from the nucleosome. Consistent with this expectation, the cryo-EM map of the ADP-BeFx-bound ncBAF-NCP complex shows an absence of the density adjacent to the acidic patch region (Figure 1A, B).

**Figure 2.**
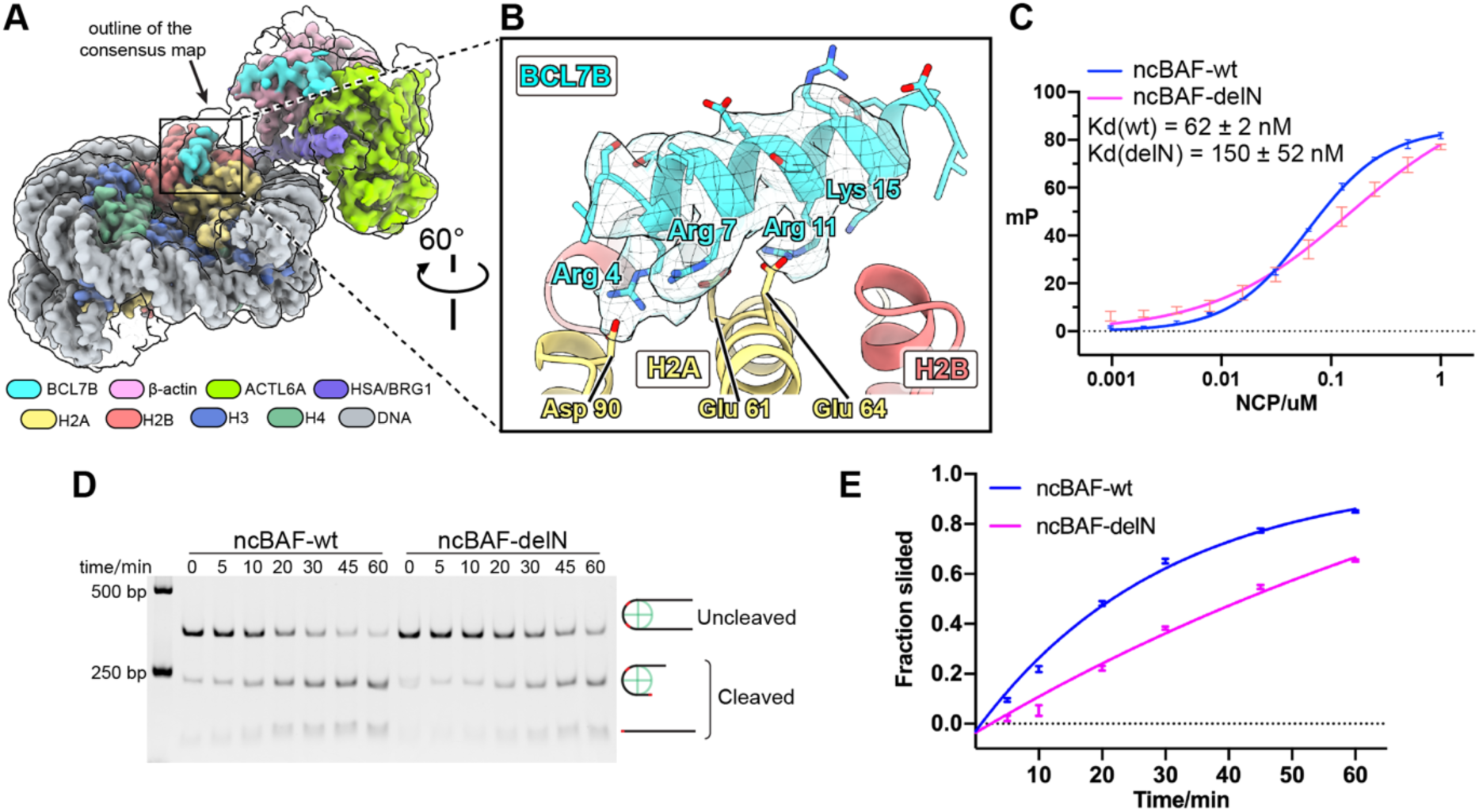
N-terminal helix of BCL7B bridges ARP and histones and contributes to nucleosome remodeling by ncBAF. (**A**) Composite cryo-EM map showing locally refined densities of the nucleosome and ARP module with the subunits colored as indicated. The overall density of the nucleosome–ARP module complex is displayed at a low contour threshold as a transparent surface, highlighting the continuous density of BCL7B extending from the ARP module toward the nucleosomal acidic patch. (**B**) Close-up view of the N-terminal helix (residues 2-18) of BCL7B engaging the nucleosomal acidic patch. The cryo-EM density of the BCL7B helix, sharpened using EMReady software, is displayed. Key basic residues of BCL7B and acidic residues of histones involved in the interaction are labeled accordingly. (**C**) Affinity measurement of the ncBAF complex for the nucleosome using fluorescence polarization binding assays. ncBAF-delN denotes the mutant complex lacking the residues 1-20 of BCL7B. The Kd values are presented as mean ±SEM, calculated from *n* = 3 independent experiments. (**D**) Restriction enzyme accessibility assay to evaluate the effect of BCL7B N-terminal helix on ncBAF-mediated nucleosome remodeling using a 100N100 nucleosome substrate. The experiment was independently repeated three times, and a representative gel image was presented. (**E**) Quantification of remodeled nucleosome fractions in panel (D) from three independent experiments using ImageJ software. Data are presented as mean ± SEM.

To examine the contribution of the N-terminal helix of BCL7B to binding the nucleosome, we used wild-type and deletion mutant (BCL7B 1-20 residues were deleted, referred to as delN) ncBAF complexes to conduct fluorescence polarization assays to measure the affinity for nucleosomes. The results showed that the deletion of the N-terminal helix in BCL7B significantly reduced the binding affinity of the apo ncBAF complex to nucleosomes by more than 50% (Figure 2C), further supporting the idea that this helix originates from the BCL7B subunit. Fluorescence polarization fitting yielded Hill coefficients of 1.216 for the wild type and 0.685 for the delN mutant. The modestly elevated coefficient of the wild type indicates weak positive cooperativity, consistent with a multivalent binding mode in which ncBAF concurrently engages two acidic patches via the BRG1 SnAC domain and BCL7B and nucleosomal DNA through the BRG1 ATPase motor. This configuration likely enables mutually reinforcing interactions between SnAC and BCL7B. In contrast, the delN mutant, which interacts with only one acidic patch, shows a Hill coefficient below unity, reflecting the loss of cooperativity and the presence of binding-site heterogeneity. This heterogeneity likely results from the asymmetric engagement of the nucleosome at a time, as only one DNA end was labeled with FITC. We further performed a nucleosome remodeling assay to evaluate the role of BCL7B-histone interactions (Figure 2D). The results demonstrated that deletion of the BCL7B N-terminal helix significantly attenuated the nucleosome sliding activity of ncBAF (Figure 2E), highlighting its essential role in ncBAF function.

Taken together, we propose that BCL7B protein can connect the ARP module to the nucleosomal acidic patch in the apo state of the ncBAF-NCP complex, and dissociate from the nucleosomes upon BRG1 binding ATP, and its interaction with the nucleosome positively contributes to ncBAF-mediated chromatin remodeling.

### Cryo-EM structures of the BCL7B-containing ARP module

The aforementioned observations suggest that the uncharacterized density within the ARP module is likely attributable to the BCL7B protein, serving as a component of the ARP module. Indeed, cross-linking mass spectrometry data of endogenous human BAF complexes showed that BCL7 proteins likely interact with the subunits of the ARP module (9). To validate this speculation and interrogate the molecular basis of BCL7 proteins in the context of the ARP module, we co-expressed human ACTL6A, β-actin, and the HSA^BRG1^ domain with BCL7B in HEK293F cells and successfully purified a four-subunit protein complex (Figure S1C). Therefore, we demonstrated that BCL7 proteins are a constitutive component of the ARP module.

To elucidate the assembly mechanism of BCL7 proteins within the ARP module, we first resolved the cryo-EM structures of the tetrameric complex in its apo (free of ATP or ADP in the sample buffer) state, resulting in a nominal resolution of 2.92 Å (Figure S7A). The high resolution allowed us to precisely model the ACTL6A, β-actin and HSA^BRG1^ subunits (Figure 3A, B). ACTL6A dimerizes with β-actin, and both subunits sequentially dock onto the HSA^BRG1^ helix in an N-to-C direction (Figure 3A, B), consistent with the architecture of their yeast counterparts (13,15). In addition to these three subunits, the high resolution of our cryo-EM map enabled an unprecedented level of detail in visualizing the integration of BCL7B within the ARP module and allowed us to unambiguously assign the remaining density adjacent to ACTL6A and β-actin to the residues from Glu26 to Val51 of BCL7B under the guiding of the distinctive densities of aromatic residues (Figure S8D, E), thereby completing the central structure of BCL7B that bridges the ARP module and the nucleosome. The unresolved C-terminal region (residues 52-end, Figure S8A) of BCL7B is unlikely to directly interact with β-actin, ACTL6A, or the HSA domain. Interestingly, both ACTL6A and β-actin contain an ATP-Mg^2+^ pair (Figure 3C, D), suggesting β-actin adopts a polymerization-competent state (41). This observation contrasts with earlier studies of the yeast ARP, where the β-actin counterpart remains a nucleotide-free pocket, while the ACTL6A counterpart tightly binds to the ATP molecule (13,15).

**Figure 3.**
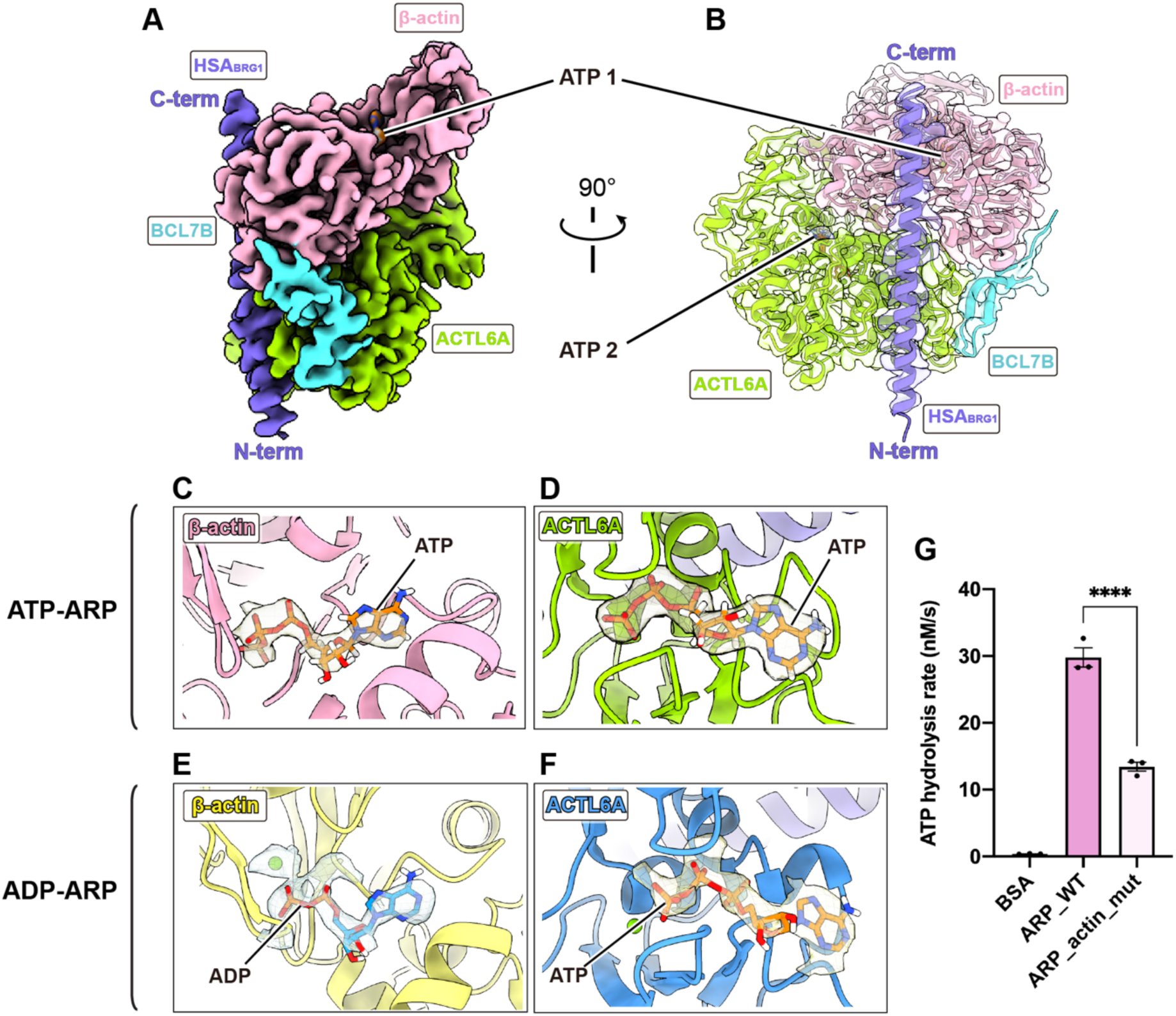
Overall structure of the BCL7B-containing ARP module. (**A**) Cryo-EM map of the ARP module in the apo state, with individual subunits colored and labeled as indicated. (**B**) Ribbon model of the ARP structure is viewed from an alternative angle, with subunits colored consistently with panel (A). The cryo-EM density map is overlaid as a transparent surface. (**C, D**) Close-up views of the nucleotide-binding pockets of the β-actin and ACTL6A in the apo ARP complexes (ATP-ARP). (**E, F**) Close-up views of the nucleotide-binding pockets of the β-actin and ACTL6A in the ADP-soaked ARP complexes (ADP-ARP). The corresponding cryo-EM densities of the bound nucleotides are shown as transparent surfaces. ADP soaking replaces the ATP molecule bound to β-actin with ADP. (**G**) Comparison of ATP hydrolysis activities between the wild-type ARP module and the actin(Q137A) mutant complex. Data are presented as mean ± SEM from three independent experiments. Statistical significance was determined by one-way ANOVA followed by Tukey’s multiple-comparison post hoc test (****p < 0.0001).

To investigate the dynamic interplay of nucleotide in ACTL6A and β-actin, we incubated the human ARP complex with ADP molecules at 4°C overnight, followed by cryo-EM single-particle analysis. The resulting map achieved a resolution of 2.74 Å, which was comparable to the apo structure (denoted as ATP-ARP because of ATP-bound β-actin) (Figure S7B). Superimposing the ATP-ARP and ADP-soaked structure (ADP-ARP) structures revealed a high structural similarity, with a root-mean-square deviation (RMSD) of 0.67 Å (Figure S9A). In the ADP-ARP structure, while ACTL6A retained an ATP-Mg²⁺ pair, β-actin transitioned to an ADP-Mg²⁺ binding state (Figure 3E, F), indicating the ATP hydrolysis potential of β-actin in the human ARP module. This nucleotide exchange in β-actin is consistent with a previous biochemical study in which β-actin accounts for approximately 1% ATP hydrolysis activity of the whole BAF complex (42). To validate β-actin’s ATPase property within the human ARP module, we co-expressed an ATP hydrolysis-deficient β-actin mutant (Q137A) (43) together with other subunits of the ARP module in HEK293F cells. The purified mutant and wild-type ARP modules were subjected to ATP-hydrolysis activity assays. The wild-type ARP module displayed weak ATPase activity, whereas the mutant complex exhibited a significantly reduced hydrolysis rate (Figure 3G). The residual ATP hydrolysis signal observed for the mutant ARP module was likely due to sample heterogeneity arising from partial incorporation of endogenous β-actin in HEK293F cells. These structural and biochemical results demonstrate that β-actin within the ARP module retains intrinsic ATP hydrolysis activity.

### Structural features of BCL7B within the ARP module

The mammalian BCL7 family proteins feature a highly conserved N-terminal region comprising the first 51 amino acids, whereas their remaining sequences show extensive variability (Figure S8A). As described above, residues 2-18 within this conserved region form an α-helix that mediates interaction with the nucleosome core (Figure 2). In the ATP-ARP structure, the residues 26-51 fold into an anti-parallel β-sheet conformation, strengthening the association between ACTL6A and β-actin by spanning across the two subunits. In the ADP-ARP structure, although the overall resolution is slightly higher, the densities corresponding to BCL7B 26-29 residues become invisible (Figure S8D, E). This loss of density is likely caused by β-actin undergoing a nucleotide transition-driven conformational change that perturbs the local environment and gives a higher flexibility for the N-terminal helix of BCL7B. Specifically, BCL7B’s Trp35 and Phe46 contribute to a hydrophobic cluster with ACTL6A’s Leu117 and Pro147, while its Trp31 and Trp48 participate in hydrophobic interactions with β-actin’s Pro367. Moreover, BCL7B residues Lys30, Trp31, Lys33, and Ser42 form hydrogen bonds and salt bridges with β-actin’s Glu125 and Glu117, as well as ACTL6A’s Ser412, respectively (Figure 4A). These multifaceted interactions collectively promote the incorporation of BCL7B and preserve the functional integrity of the ARP module.

**Figure 4.**
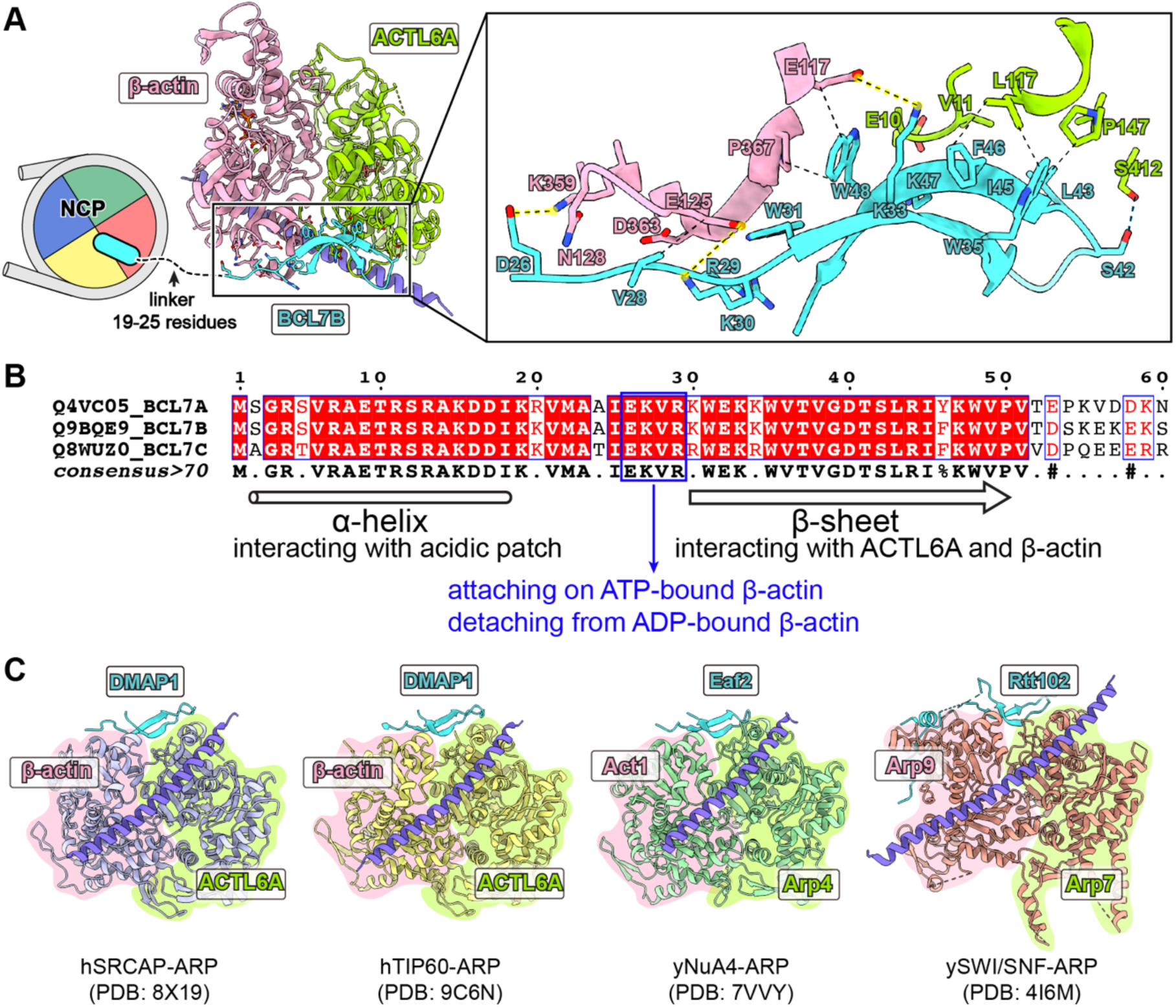
BCL7B offers a conserved β-sheet to stabilize the ARP module. (**A**) ATP-ARP structure and the close-up view of the BCL7B subunit. The N-terminal helix of BCL7B bound to the nucleosome is shown in cartoon representation. Key residues involved in the assembly of BCL7B into the ARP module are labeled and displayed in stick mode. Salt bridges, hydrophobic interactions, and hydrogen bonds formed between the BCL7B and β-actin/ACTL6A are illustrated using dashed lines, with salt bridges highlighted in yellow shading. (**B**) Multiple sequence alignment of the N-terminal region (1-60 residues) in BCL7 family proteins with the conserved residues highlighted in red blocks. Secondary structure elements resolved in our apo ARP structure are annotated as indicated. Residues 26-29, which lose cryo-EM density in the ADP-ARP complex, are outlined in blue, indicating subtly increased local flexibility of BCL7B and structural rearrangement upon nucleotide exchange of β-actin. (**C**) Local structures of the ARP module in other chromatin-associated complexes, including human SRCAP and TIP60, as well as yeast NuA4 and SWI/SNF. PDB IDs of the corresponding structures are annotated in the panel. The conserved β-sheets derived from non-homologous proteins are shown in cyan and the HSA domains are in slate.

Despite a lack of sequence similarity, the anti-parallel β-sheet structure of BCL7B aligns pretty well with the β-sheet of Rtt102 in yeast ARP, with an RMSD of 0.27 Å (Figure S8F). In a broader chromatin regulation complex context, recent cryo-EM reconstructions of the human chromatin remodeling SRCAP complex bound to nucleosomes, along with the TIP60 complex reveal analogous assembly patterns of the ARP module (16–19). In these structures, the corresponding β-sheet originates from the DMAP1 subunit (Figure 4C). This conserved structural motif is also present in the yeast NuA4 subunit Eaf2, where a similar β-sheet spans the interface between Act1 and Arp4 (20). Collectively, these structural observations elucidate the assembly mechanism of BCL7B within the ARP module and reveal a conserved β-sheet fold across diverse complexes and species.

### Structural features of ACTL6A and β-actin in the ARP module

Enhanced incorporation of ACTL6A into the SWI/SNF complex has been linked to cancer progression (44,45), and the switching of ACLT6A/B orthologs within the complex is essential for neural development (46–48). These observations underscore the critical role of ACTL6A/B proteins in SWI/SNF-mediated chromatin remodeling. As an actin-related protein, ACTL6A/B exhibits distinct functional properties from β-actin, most notably lacking ATP hydrolysis activity and the ability to polymerize (49). Our high-resolution cryo-EM structures of the ARP module offer critical insights into the structural mechanisms that underlie these functional differences. Although ACTL6A exhibits substantial structural similarity to β-actin (RMSD 1.57 Å, Figure 5A), a notable difference lies in an insertion structure (Met213-Gln245, Figure 5A). β-actin filament elongation depends on the combined interface of subdomain 2 (SD2) and SD4 (Figure 5A, Figure S10B) (50). When ACTL6A is aligned into the β-actin filament, the insertion occludes the SD4 interface and overlaps with the adjacent β-actin subunit, which creates a clear steric hindrance to prevent ACTL6 from undergoing a β-actin-like polymerization (Figure 5A, Figure S10C). The D-loop region (Val45-Gly65) of ACTL6A remains unresolved in our structure, indicating inherent flexibility like in β-actin (Figure 5A). While ATP binding residues are largely conserved between β-actin and ACTL6A/B, two residues essential for ATP hydrolysis in β-actin, Gln137 and His161 (43), are replaced by Thr154 and Thr178 in ACTL6A/B, respectively (Figure 5B, Figure S10A). These substitutions likely abrogate ATP hydrolysis by disrupting the water bridges that are essential for the hydrolysis process (43). The absence of ATPase activity in ACTL6A/B proteins is supposed to stabilize their association with β-actin within the ARP module.

**Figure 5.**
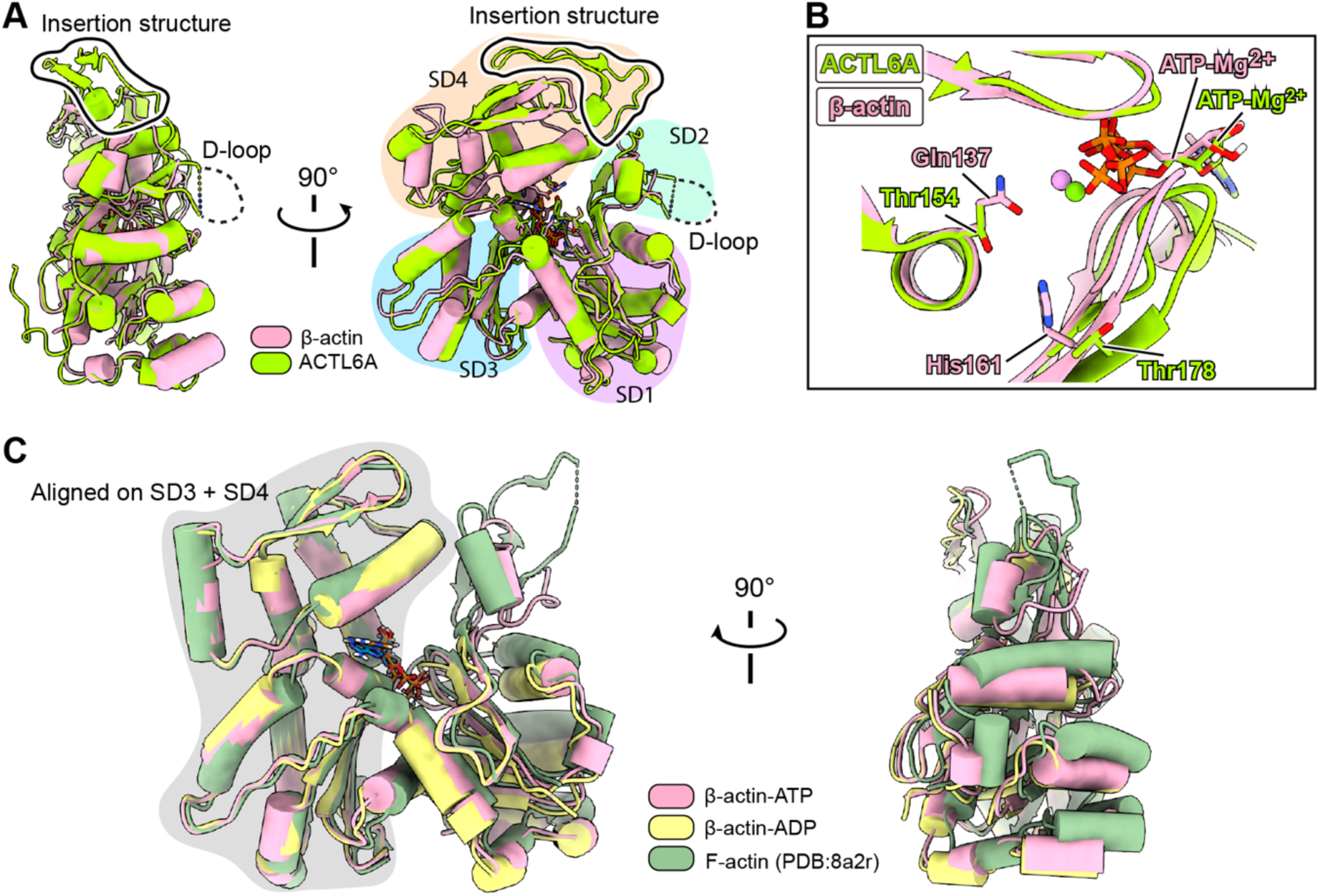
Structural features of β-actin and ACTL6A. (**A**) A superimposition of ACTL6A and β-actin in the ATP-ARP structure, with the ACTL6A-specific insertion segment outlined and unstructured D-loop annotated. The four subdomains (SD1-SD4) of β-actin are delineated by color lumps. **(B**) Close-up view of the nucleotide-binding pockets of the aligned structures in ribbon cartoon from panel (**A**), showing Gln137 and His161 of β-actin and Thr154 and Thr178 of ACTL6A in stick mode. (**C**) Three β-actin molecules, two from our ATP-ARP and ADP-ARP structures and one from an actin filament (PDB ID: 8a2r) are superimposed based on their SD3 and SD4 subdomains. Two orthogonal views of the alignment are presented.

Upon nucleotide transition from ATP to ADP following incubation of ARP module complex in an ADP-containing buffer, the SD2 subdomain of β-actin exhibited increased flexibility, resulting in poor resolution of the His73 loop (residues 50-79) and making it difficult to model (Figure S9B). Structural alignment of β-actin molecules from the ATP/ADP-ARP structures, based on their inner domains (SD3 and SD4), revealed subtle conformational shifts in their outer domains (SD1 and SD2) (Figure 5C). These observations indicate that the conformation of β-actin within the ARP module is regulated by its nucleotide-binding states. Due to the absence of induced fitting and the structural constraints imposed by the neighboring β-actin in the actin filament, the conformation of β-actin within the ATP-ARP structure deviates from that observed in filamentous actin (Figure 5C). It is worth noting that although ATP was not supplemented in the sample buffer, the retention of ATP in β-actin within the ATP-ARP structure may be attributed to its low intrinsic hydrolysis rate (41).

### ARP module structurally enables SWI/SNF interaction with F-actin barbed end

Mechanotransduction pathways critically influence chromatin organization and gene transcription, with nuclear actins playing a pivotal role in these processes (51). Interactions between ARP-containing chromatin remodelers — such as the SWI/SNF, INO80, and SWR1 complexes — and nuclear actin filaments have been observed both in vitro and in vivo (52–54), yet the precise binding mechanism remains unclear. Our combined structural and biochemical analyses of the human ARP module establish that β-actin within SWI/SNF complexes can hydrolyze ATP. Given that the ATP-bound state of β-actin is indispensable for its polymerization and filament elongation (51), we are interested in exploring whether the nucleotide binding-competent β-actin with the pointed end exposed in the SWI/SNF complexes has the potential to incorporate into the barbed end of an actin filament. Superimposition of the SWI/SNF complexes onto the terminal β-actin subunit at the barbed end leads to no steric clashes with adjacent subunits, suggesting a structurally compatible interface between the ARP module and the actin filaments (Figure 6A). AlphaFold 3 generates a high-confidence model of the ARP module bound to an actin filament (Figure 6B, C), providing further support for this potential mechanism. Indeed, the polymerization-deficient β-actin mutant (R62D, in subdomain 2) completely abolishes its in vivo interaction with the SWI/SNF complexes (52), thereby supporting the proposed interaction mode in which ARP module mediates the association of SWI/SNF complexes with actin filaments to coordinate the mechanotransduction signaling with chromatin remodeling.

**Figure 6.**
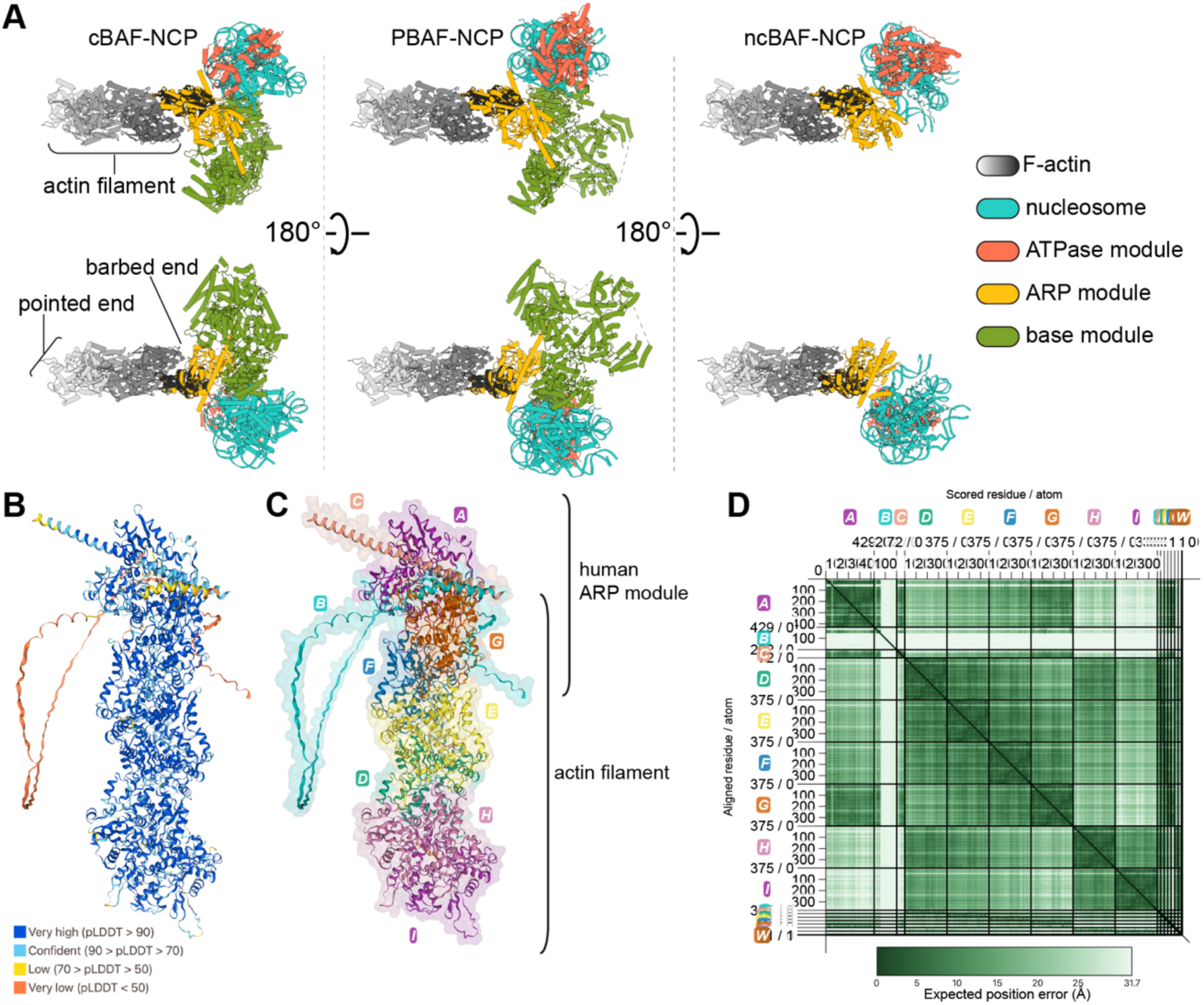
A potential interaction mode of SWI/SNF complexes with actin filaments. (**A**) Structures of cBAF (PDB: 6ltj), PBAF (PDB: 7vdv), and our apo ncBAF are superimposed onto the terminal β-actin subunit of an F-actin filament structure (PDB: 8a2r), revealing no steric clashes between the actin filament and any SWI/SNF subunits. The SWI/SNF modules are colored and labeled as indicated with the terminal β-actin in the filament particularly colored in black. (**B**) Predicted complex structure generated by the AlphaFold 3 server, using one ACTL6A, one BCL7B, one HSA^BRG1^ domain (447-518 residues), six β-actin molecules, and seven ATP-Mg^2+^ pairs as input. **c**, The predicted model from panel (**B**) colored by subunit and labeled A-I (A: ACTL6A, B: BCL7B, C: HSA^BRG1^ domain, D-I: β-actin). Notably, β-actin subunit G is shared between the ARP module and actin filament. (**D**) Predicted aligned error (PAE) plot for the predicted ARP–actin filament model in panel (**B**), illustrating the predicted positional uncertainty both within and between input items. Axes are labeled A–I for protein chains (as in panel (**C**)), J–P for ATP molecules, and Q–W for Mg²⁺ ions. Graphs in panels (**B**-**D)** are obtained from the PAE Viewer (https://subtiwiki.uni-goettingen.de/v4/paeViewerDemo) following analysis of the AlphaFold 3 prediction results.

## Discussion

Among the three SWI/SNF complex variants, ncBAF exhibits a distinct chromatin targeting profile, characterized by a preferential association with CTCF-enriched loci and promoter regions (5). Through integrated structural characterization and cross-linking mass spectrometry of the ncBAF-nucleosome complexes under ATP analog-bound and -free conditions, we identified fundamental differences in nucleosome engagement architectures between ncBAF and cBAF/PBAF. Notably, while cBAF and PBAF sandwich nucleosomes through the ATPase and base modules, ncBAF establishes a bipartite nucleosome clamp by utilizing the newly identified component BCL7B of the ARP module instead of the base module. This unique mode arises from the absence of SMARCB1 in ncBAF, which in cBAF/PBAF stabilizes the base module interaction with the nucleosomal acidic patch. Our XL-MS data further support this, showing fewer histone-base module cross-links in ncBAF compared to cBAF/PBAF. As we prepare to submit the manuscript, a paper also reported that the conserved N-terminal region of BCL7A can associate with the nucleosomal acidic patch (55). Given that BCL7 proteins are also present in cBAF and PBAF, it is intriguing to elucidate the coordinated mechanism between BCL7 and SMARCB1 in SWI/SNF-mediated chromatin remodeling in vivo. The interaction of BCL7B with histones further highlights the hub role of the ARP module, serving as a structural and functional nexus that links multiple submodules of the remodeler and connects the remodeler to its nucleosome substrate. A similar architectural principle is observed in the human TIP60 complex, an evolutionarily integrated supercomplex that combines chromatin remodeling and acyltransferase functions (18,19).

Previous studies have shown that BCL7 depletion reduces chromatin accessibility without altering the global chromatin-targeting profile of SWI/SNF complexes (21,56), yet the mechanistic basis for this phenotype has remained unknown. Our focused structural analyses further reveal previously unappreciated conformational dynamics of the ARP module across BRG1’s ATP-hydrolysis cycle, in which the periodic association and dissociation of BCL7B-histone are driven by nucleotide-dependent rearrangements of the ARP module. This nucleotide-responsive interaction provides a regulatory mechanism that fine-tunes nucleosome engagement and remodeling by ncBAF.

Unexpectedly, we demonstrate that β-actin within the ARP module retains ATP hydrolysis activity. The four BCL7B residues (26–29) that anchor to ATP-bound β-actin become invisible in the ADP-bound state, likely due to conformational changes induced by nucleotide transition (Figure 4B, 5C, Figure S8D, E). This observation raises an intriguing question about whether the nucleotide-binding states of β-actin within the ARP module influence BCL7’s role in ncBAF-mediated chromatin remodeling by modulating the flexibility of the BCL7 N-terminal helix, or conversely, whether the BCL7-histone engagement affects β-actin’s nucleotide-binding states through changes in the flexibility of the BCL7 N-terminal region.

## Data Availability

Coordinates and Cryo-EM density maps have been deposited in the Electron Microscopy Data Bank and Protein Data Bank under accession codes EMD-64578, 9UXA (ncBAF–NCP complex in apo state); EMD-64577, 9UX9 (ncBAF–NCP complex in ADP-BeFx-bound state); EMD-64579, 9UXB (ATP-ARP); and EMD-64580, 9UXC (ADP-ARP), respectively.

## Acknowledgements

We thank the cryo-EM Center, Guangzhou Institutes of Biomedicine and Health Chinese Academy of Sciences and Advanced Bio-imaging Technology Platform of Guangzhou Laboratory for cryo-EM beamtime and all staff members for their assistance with data collection. We also thank the Mass Spectrometry System staff members at the National Facility for Protein Science in Shanghai (NFPS), Zhangjiang Lab, China for technical support and assistance in XL-MS data collection and analysis.

## Author contributions

J.H., Y.Z. and F.S. designed the project. F.S. and P.X. constructed the expression plasmids for ncBAF. F.S., Q.L. and C.W. prepared protein complexes and the nucleosomes involved in the research. F.S. and B.Z. conducted cryoEM experiments and determined the structures. F.S. conducted the biochemical experiments. B.Z., F.S. and H.L. prepared the structures and figures under the supervision of D.Q. and J.H.. C.X. and J.C. discussed the results and provided functional feedback in the manuscript. F.S. and J.H. prepared the manuscript with input from all authors. J.H. Y.Z. and D.Q. supervised and directed the overall research.

